# Spatiotemporal specification of human green and red cones

**DOI:** 10.1101/2021.03.30.437763

**Authors:** Sarah E. Hadyniak, Kiara C. Eldred, Boris Brenerman, Katarzyna A. Hussey, Joanna F. D. Hagen, Rajiv C. McCoy, Michael E. G. Sauria, James A. Kuchenbecker, Thomas Reh, Ian Glass, Maureen Neitz, Jay Neitz, James Taylor, Robert J. Johnston

## Abstract

Trichromacy is unique to primates among mammals, enabled by blue (short/S), green (medium/M), and red (long/L) cones. In humans and Old World monkeys, cones make a poorly understood choice between M and L cone subtype fates. To determine mechanisms specifying M and L cones, we developed an approach to visualize expression of the highly similar *M*- and *L*-*opsin* mRNAs. *M*-*opsin*, but not *L*-*opsin*, was observed during early human eye development, suggesting that M cones are generated before L cones. In adult human tissue, the early-developing central retina contained a mix of M and L cones compared to the late-developing peripheral region, which contained a high proportion of L cones. Retinoic acid (RA)-synthesizing enzymes are highly expressed early in retinal development. High RA signaling early was sufficient to promote M cone fate and suppress L cone fate in retinal organoids. Across a human population sample, natural variation in the ratios of M and L cone subtypes was associated with a noncoding polymorphism in the *NR2F2* gene, a mediator of RA signaling. Our data suggest that RA promotes M cone fate early in development to generate the pattern of M and L cones across the human retina.

## Introduction

Trichromacy in humans is enabled by three subtypes of cone photoreceptors that express opsin photopigments sensitive to short (S), medium (M), or long (L) wavelengths of light. Among mammals, only humans, Old World primates, and some New World monkeys possess M and L cones (*1*–*3*). The only known difference between M and L cones is the expression of their opsin photopigment, OPN1MW (M-opsin) or OPN1LW (L-opsin) (*4*). M and L cones have been technically challenging to distinguish due to the high sequence and structural similarity between the M- and L-opsin proteins, and thus, the mechanisms underlying this fate decision have proven elusive.

In humans, the *OPN1LW* (*L-opsin*) and *OPN1MW* (*M-opsin*) genes lie in a tandem array on the X chromosome under the control of a shared regulatory DNA element called a locus control region (LCR) (*5*–*8*). Though the number and ordering of *OPN1LW* and *OPN1MW* genes varies among individuals, the LCR is most commonly found upstream of *OPN1LW* followed by two copies of *OPN1MW* (*9*). Regardless of the arrangement, only the first two opsin genes in the array are expressed (*4*, *5*).

Two nonexclusive mechanisms have been proposed to control M and L cone fates. The stochastic model, which suggests that cones randomly choose M or L cone fate, is based on transgene reporter experiments in mice (*6*, *10*). The temporal model is supported by functional microspectrophotometry, multifocal electroretinograms (ERGs), and qPCR-based expression studies, which found that the late-born retinal periphery is enriched for L cones compared to the early-born central retina (*11*–*13*). In this study, we examined human retinas and manipulated retinal organoids to assess these mechanisms.

Organoids are powerful systems to investigate the molecular mechanisms controlling cell fate specification during human development (*14*). We previously showed that thyroid hormone regulates the S vs. M/L cone fate decision in human retinal organoids (*15*). Here, we studied the development of human retinas and organoids and conducted an association study to understand the M vs. L cone fate decision. Our data suggest that retinoic acid (RA) signaling controls the spatiotemporal patterning of M and L cones in the developing human retina.

## Results

### Visualization of *M*- and *L*-*opsin* expression

To observe cone fates, we developed a method to visualize *M*- and *L*-*opsin* mRNA expression. The opsin protein sequences are ~96% identical and the mRNA sequences are ~98% identical, complicating expression analysis. To directly visualize *M*- and *L*-*opsin* expression, we generated a 40-nucleotide probe for *M*-*opsin* and a 42-nucleotide probe for *L*-*opsin* with partial overlap that target a region with 8 differential nucleotides (**Fig. 1A-B**). We utilized colorimetric *in situ* hybridization to visualize *M*-*opsin* in blue and *L*-*opsin* in pink.

**Fig. 1.**
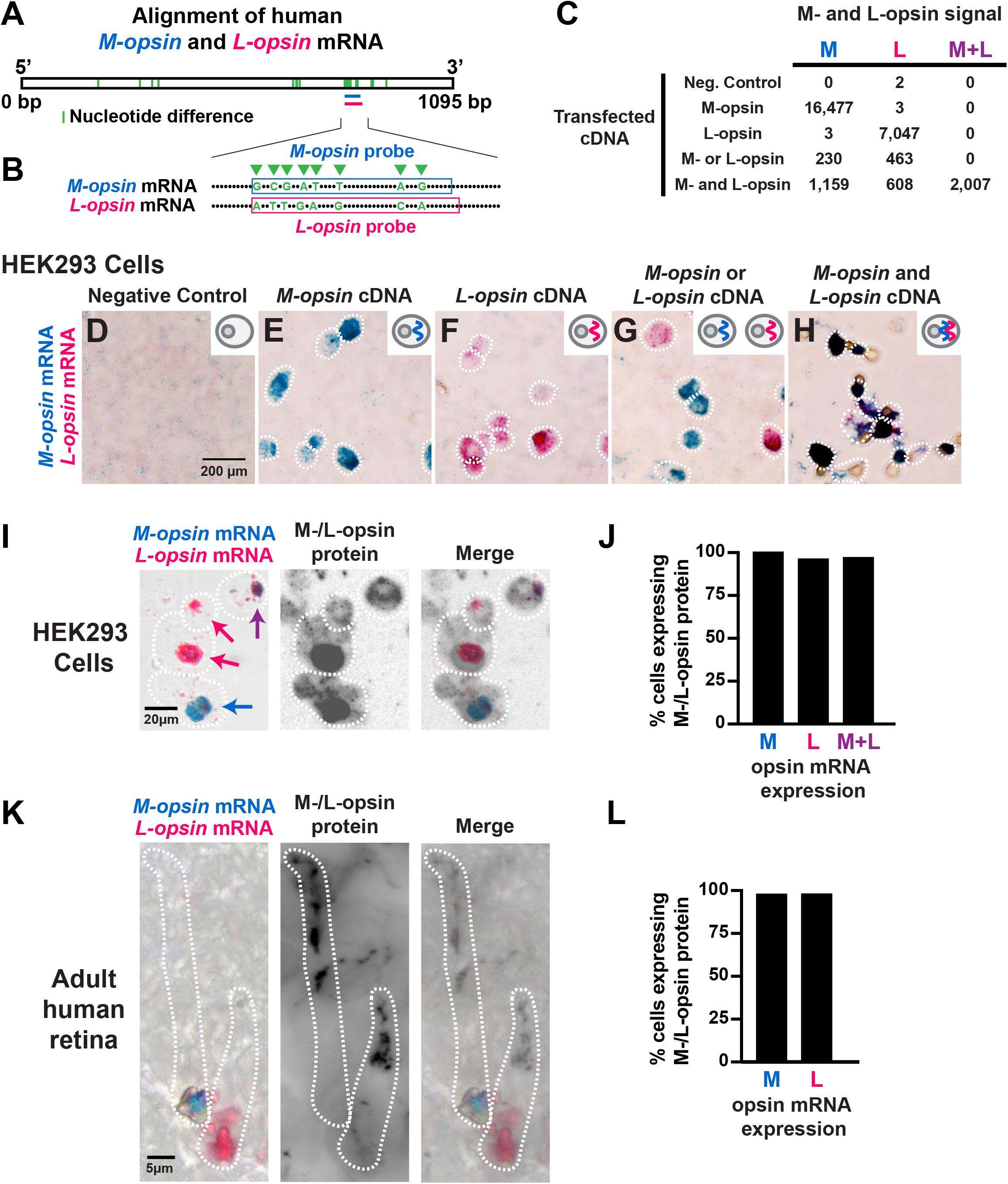
An *in situ* hybridization approach distinguishes *M*-*opsin* and *L*-*opsin* mRNA. **(A)** Alignment of human *M*-*opsin* and *L*-*opsin* mRNA. Green bar=nucleotide difference. Horizontal pink and blue lines=location of *in situ* probes. **(B)** Alignment of portions of exon 5 from *M*- and *L*-*opsin*. *in situ* hybridization probe binding sites are indicated by blue (*M*-*opsin*) and pink (*L*-*opsin*) boxes. Green arrowheads indicate 8 nucleotide differences. Dots indicate nucleotide alignment between the opsins. **(C-H)** HEK293 cells probed for *M*-*opsin* mRNA (blue) and *L*-*opsin* mRNA (pink). Insets=schematic of transfected plasmid. Cells that did not express *M*-*opsin* mRNA and/or *L*-*opsin* mRNA were not quantified. **(C)** Quantification of cells expressing *M*-*opsin* mRNA only, *L*-*opsin* mRNA only, or *M*-*opsin* mRNA and *L*-*opsin* mRNA for the conditions in (**D-H**). **(D)** No plasmid transfected. **(E)** Transfection of a plasmid driving *M*-*opsin*. **(F)** Transfection of a plasmid driving *L*-*opsin*. **(G)** Transfection of either a plasmid driving *M*-*opsin* or a plasmid driving *L*-*opsin* independently and then the cells were mixed. **(H)** Transfection of both a plasmid driving *M*-*opsin* and a plasmid driving *L*-*opsin*. **(I)** Visualization of *M*-*opsin* mRNA, *L*-*opsin* mRNA, and M-/L-opsin protein in HEK293 cells transfected with both a plasmid driving *M*-*opsin* and a plasmid driving *L*-*opsin*. *M*-*opsin* (blue) and *L*-*opsin* (pink). Blue arrow indicates a cell expressing *M*-*opsin* mRNA only. Pink arrows indicate cell expressing *L*-*opsin* mRNA only. Purple arrow indicates a cell expressing both *M*-*opsin* mRNA and *L*-*opsin* mRNA. **(J)** Quantification of **(I)**. **(K)** Visualization of *M*-*opsin* mRNA, *L*-*opsin* mRNA, and M-/L-opsin protein in cone cells in an adult human retina. *M*-*opsin* (blue) and *L*-*opsin* (pink). No cones coexpressed *M*-*opsin* mRNA and *L*-*opsin* mRNA. **(L)** Quantification of **(K)**.

We tested the specificity of this approach to distinguish *M*- and *L*-*opsin* mRNA by transfecting plasmids driving *M*-*opsin* and/or *L*-*opsin* and visualizing expression in HEK293 cells. We scored the total number of transfected cells that expressed *M*-*opsin* mRNA (**Fig. 1C**, “M”), *L*-*opsin* mRNA (**Fig. 1C**, “L”), or both *M*-*opsin* and *L*-*opsin* mRNA (**Fig. 1C**, “M+L”) for each experiment. HEK293 cells transfected with no plasmid showed almost no expression (**Fig. 1C-D**). Cells transfected with a plasmid driving *M*-*opsin* showed expression of *M*-*opsin* but not *L*-*opsin* (**Fig. 1C, 1E**), while cells transfected with a plasmid driving *L*-*opsin* displayed expression of *L*-*opsin* but not *M*-*opsin* (**Fig. 1C, 1F**). Cells transfected with either a plasmid driving *M*-*opsin* or a plasmid driving *L*-*opsin* and then mixed, showed expression of *M*-*opsin* only or *L*-*opsin* only in individual cells (**Fig. 1C, 1G**). Finally, cells co-transfected with both a plasmid driving *M*-*opsin* and a plasmid driving *L*-*opsin* displayed co-expression of *M*- and *L*-*opsin* (**Fig. 1C, 1H**). In this experiment, cells expressing *M*-*opsin* or *L*-*opsin* only were also observed, due to variation in transfection of the two plasmids (**Fig. 1C**). Thus, this *in situ* hybridization method distinguishes between the highly similar *M*- and *L*-*opsin* mRNAs.

We next related expression of opsin mRNA and protein. Due to the high similarity of the M-opsin and L-opsin proteins, available antibodies detect both proteins. We transfected HEK293 cells with both a plasmid driving *M*-*opsin* and a plasmid driving *L*-*opsin* and observed cells that expressed *M*-*opsin* only, *L*-*opsin* only, or both *M*-*opsin* and *L*-*opsin* (**Fig. 1I**), like our previous experiment (**Fig. 1C**). We examined protein expression with an M-/L-opsin antibody and observed expression of M-/L-opsin protein in nearly all cells that expressed *M*-*opsin* alone, *L*-*opsin* alone, or both *M*-*opsin* and *L*-*opsin* (**Fig. 1I-J**), showing that opsin mRNA-expressing cells identified by this method also express opsin protein.

We next tested this assay in adult human retinas. We identified cells that exclusively expressed *M*-*opsin* or *L*-*opsin* (**Fig. 1K**). M-/L-opsin protein was observed in nearly all cells that expressed *M*-*opsin* or *L*-*opsin* mRNA (**Fig. 1K-L**), suggesting that this method identifies opsin protein-expressing M- and L-cones.

Together, we conclude that this *in situ* hybridization method distinguishes between *M*- and *L*-*opsin* mRNAs and reliably identifies M-cones and L-cones in adult human retinas.

### *M*-*opsin* is expressed before *L*-*opsin* during development

To assess the timing of M and L cone specification during development, we evaluated *M*-*opsin* or *L*-*opsin* mRNA in a 122-day old human fetal retina. We predicted two main, alternate outcomes: (1) a mix of cones expressing either *M*-*opsin* or *L*-*opsin* if the decision was stochastic, or (2) one population of cones expressing *M*-*opsin* or *L*-*opsin* only if the decision was temporal and one cone subtype was generated first. In this midgestation retina, we observed *M*-*opsin* and no *L*-*opsin* (100% *M*-*opsin* only cells, n>500 cells) (**Fig. 2A**). This retina displayed M-/L-opsin protein (**Fig. S1A**), suggesting that these cells were terminally differentiating M cones. This retina also displayed Ki67 expression (**Fig. S1A**), suggesting that there are proliferating cells, ongoing development, and generation of additional neurons at this timepoint in this region of the fetal retina. Together, these data suggested that M cones are generated before L cones during human retinal development.

**Fig. 2.**
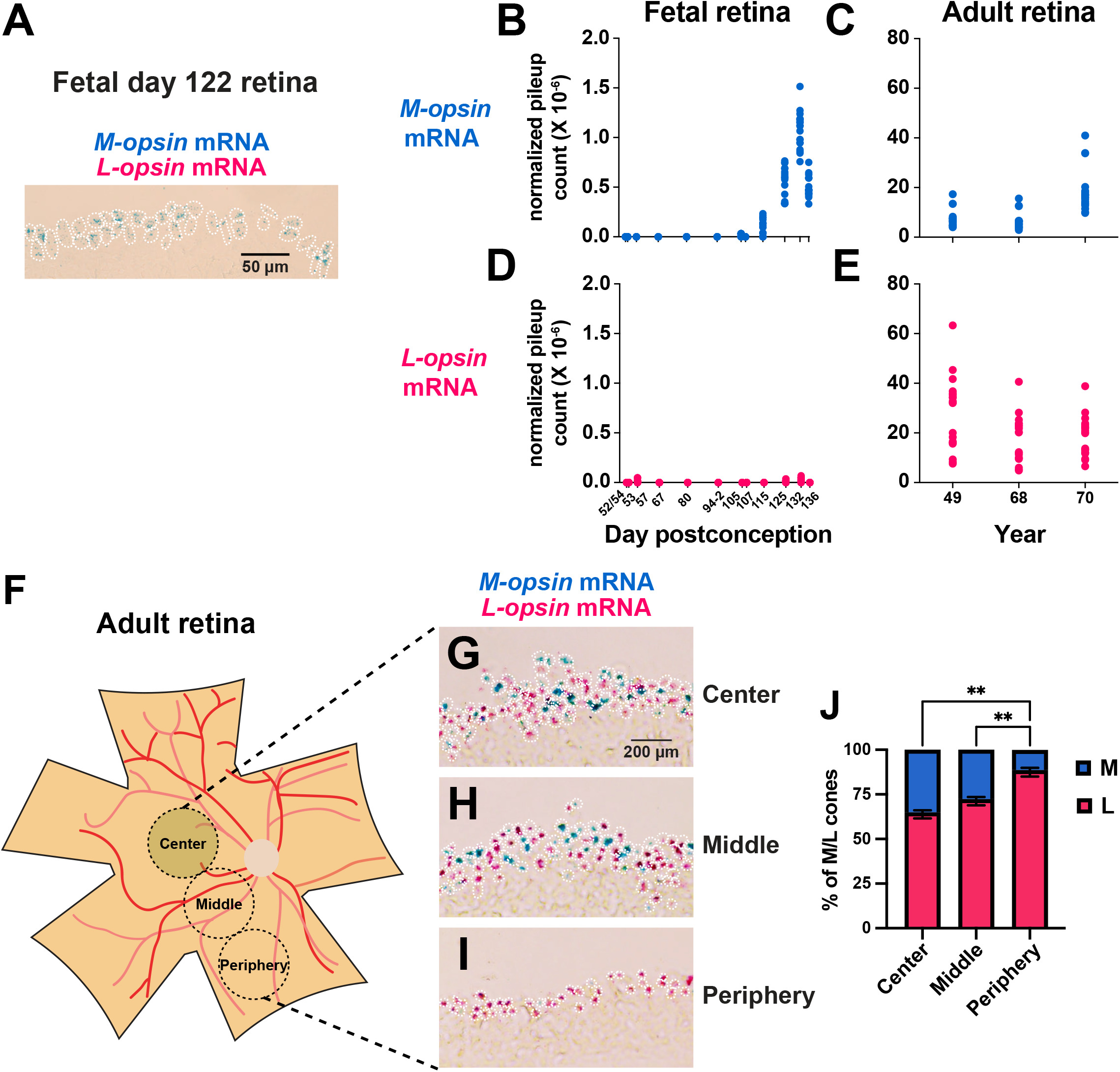
Expression of *M*-*opsin* mRNA and *L*-*opsin* mRNA in fetal and adult human retinas. **(A)** 20 μm section of a fetal day 122 retina. *M*-*opsin* (blue) and *L*-*opsin* (pink) **(B-E)** *M*- and *L*-*opsin* expression. Values indicate total pileup count normalized to total read count. Each data point indicates one nucleotide difference in *M*- or *L*-*opsin*. N=1 for each timepoint, except 94-2 where N=2. 52/54=exact date is unclear. Analyzed from (*16*, *17*). **(B)** *M*-*opsin* mRNA in fetal retinas. **(C)** *M*-*opsin* mRNA in adult retinas. **(D)** *L*-*opsin* mRNA in fetal retinas. **(E)** *L*-*opsin* mRNA in adult retinas. **(F)** Schematic of retina with regions isolated using a 5 mm biopsy punch. White circle=optic nerve. Red lines=blood vessels. Yellow circle = macular pigment. **(G-I)** 20 μm sections were probed for *M*-*opsin* (blue) and *L*-*opsin* (pink) mRNA. **(J)** Average ratios of M and L cones as percent of M/L total cones across three individuals. One-way ANOVA with Tukey’s multiple comparisons test: Center L versus Middle L=no significance; Center L versus Periphery L p<0.01; Middle L versus Periphery L p<0.01.

We next quantified *M*- and *L*-*opsin* expression in RNA-seq datasets by comparing counts of aligned sequencing reads for *M*- or *L*-*opsin* based on 20 nucleotide differences that distinguish these genes (**Fig. 1A**), normalizing by the total number of aligned reads per sample (“normalized pileup count”). For these experiments, each point represents the normalized pileup count for reads with one nucleotide difference for *M*-*opsin* mRNA or *L*-*opsin* mRNA in that individual sample.

We validated this approach in the Weri-Rb-1 retinoblastoma cell line, which expresses both *M*-*opsin* and *L*-*opsin* upon addition of thyroid hormone (T3) (**Fig. S1B**). We evaluated expression in 12 prenatal samples from day 52/54 through day 136 postconception from published RNA-seq datasets (*16*). *M*-*opsin* expression was observed from day 115 through day 136 (**Fig. 2B**). In contrast, *L*-*opsin* expression was not detected during this time (**Fig. 2D**). We analyzed expression from three independent adult human retinas from published datasets (*17*) and observed expression of both *M*- and *L*-*opsin* (**Fig. 2C, 2E**). These data suggest that *M*-*opsin* is expressed before *L*-*opsin* during human development.

To assess cone distributions in the adult human retina, we dissected three retinas, isolated and sectioned tissue from the center, middle, and periphery, and performed our *in situ* hybridization strategy. The periphery had a significantly higher proportion of L cones compared to the center and middle (**Fig. 2F-J; S1C-E**). As cone specification occurs from the center to the periphery during development (*18*, *19*), the higher proportions of M cones in the earlier-specified center and middle regions are consistent with the generation of M cones before L cones.

### Early retinoic acid signaling promotes M cones and suppresses L cones in human retinal organoids

We next sought to identify a mechanism that regulates M and L cone specification. We assessed expression of RA pathway regulatory genes during early human retinal development in a published dataset (*16*). Specific aldehyde dehydrogenases (ALDHs) convert retinaldehyde to RA. *ALDH1A1* and *ALDH1A3* are expressed early in retinal development and decrease over time, whereas *ALDH1A2* is expressed at very low levels throughout gestation (**Fig. 3A**). Similar to *ALDH1A1 and ALDH1A3* in humans (**Fig. 3A**), *ALDH* orthologs in mouse (*20*), zebrafish (*21*), and chicken (*22*), are expressed highly early and decrease during development. Cytochrome p450 family 26 (CYP26) enzymes catalyze the degradation of RA. *CYP26A1* is steadily expressed at low levels, whereas *CYP26B1* and *CYP26C1* are not expressed (**Fig. S2A**). The high expression of two RA-synthesizing enzymes early suggested that RA signaling is high early and decreases during development. As *M*-*opsin* is expressed before *L*-*opsin*, we hypothesized that high RA signaling early promotes M cone fate and suppresses L cone fate.

**Fig. 3.**
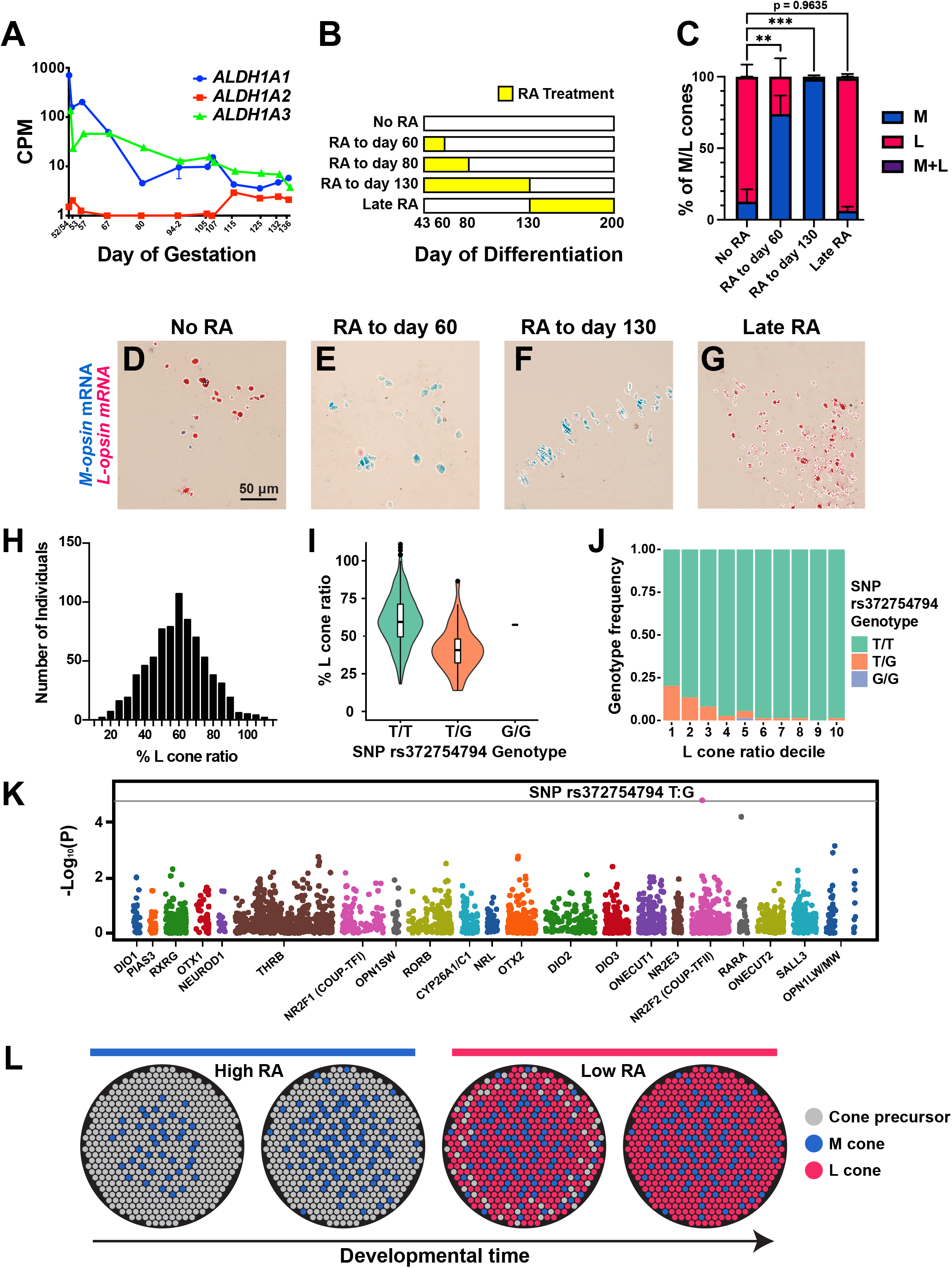
RA signaling induces *M*-*opsin* and inhibits *L*-*opsin* early in human retinal organoids. **(A)** Expression of *ALDH1A1, ALDH1A2*, and *ALDH1A3* in fetal human retinas. CPM=log counts per million. Analyzed from (*16*). **(B)** Yellow bars indicate temporal windows of 1.0 μM RA addition. **(C)** Quantification of M and L cone ratios for RA treatments. For ‘No RA’, N=3; for ‘RA to day 60’, N=6; for ‘RA to day 130’, N=3; and for ‘Late RA’, N=5. One-way ANOVA with Dunnett’s multiple comparisons test: ‘No RA’ *L*-*opsin* versus ‘RA to day 60’ *L*-*opsin*, p<0.05; ‘No RA’ *L*-*opsin* versus ‘RA to day 130’ *L*-*opsin* p<0.005; ‘No RA’ *L*-*opsin* versus ‘Late RA’ *L*-*opsin* p=0.994. Error bars indicate SEM. **(D-G)** *M*-*opsin* (blue) and *L*-*opsin* (pink) expression in organoids grown in different RA conditions (**B**), quantified in (**C**). White outlines indicate *M*- or *L*-*opsin*-expressing cells. **(D)** No RA. **(E)** RA to day 60. **(F)** RA to day 130. **(G)** Late RA. **(H)** Histogram of the ratios of % L from 738 human males with normal color vision. **(I)** L cone ratios for SNP rs372754794 genotypes. **(J)** SNP rs372754794 genotype frequencies for L cone ratio deciles. **(K)** Manhattan plot of the genetic variant p-values. Genetic variants above the gray line (Bonferroni corrected threshold) are significant (p<0.05). **(L)** Model: RA regulates the spatiotemporal patterning of M and L cones in the human retina.

We tested this hypothesis using human retinal organoids. Differentiation of human retinal organoids involves addition of 1.0 μM RA on days 20-43 to promote early retinal patterning (**Fig. S2B**). Organoids were grown in media without exogenous RA from day 43 to day 200, from the end of primitive retina differentiation through early cell fate specification (**Fig. 3B, ‘No RA’**). These organoids displayed a mix of M cones and L cones (**Fig. 3C-D**), similar to adult human retinas (**Fig. 2G-J**).

To test whether RA was sufficient to promote M cone specification in retinal organoids, we added 1.0 μM RA over different timeframes and assessed M and L cone ratios at day 200 (**Fig. 3B**). Organoids grown in supplemental RA throughout development failed to normally differentiate, yielding minimal to no M or L cones (N=6 experiments, data not shown). We previously examined gene expression using bulk RNA-seq at multiple timepoints during organoid development (*15*). We analyzed this data and found that expression of *THRB*, a marker of cone fate, stabilizes on day 70 (**Fig. S2C),** suggesting that the generation of new cones ends around this timepoint. Opsin expression was first observed at day 130 (*15*), suggesting that terminal differentiation of cones begins at this timepoint. We defined three main timeframes of cone development: immature cone generation through day 70, maturation from days 70 to 130, and terminal differentiation from day 130 onwards.

Addition of RA early from days 43 to 130 yielded organoids with almost exclusively M cones at day 200 (98% M, 2% L, 0% co-expressing; **Fig. 3C, 3F**; **‘RA to day 130’**), suggesting that addition of RA through the beginning of *M*- and *L*-*opsin* expression was sufficient to promote M cone fate. To determine whether RA could promote M cone fate in immature cones, we added RA through day 60, before the end of immature cone generation and well before terminal differentiation and *M*- and *L*-*opsin* expression on day 130. In this condition, organoids were enriched for M cones (74% M, 26% L; **Fig. 3C, 3E;** ‘**RA to day 60**’), suggesting that RA was sufficient to promote M cone fate in immature cones. Addition of RA late in development from days 130 to 200 yielded L cone-enriched organoids at day 200 (6% M, 93% L, 1% co-expressing; **Fig. 3C, 3G; ‘Late RA’**), similar to ‘No RA’ control organoids (**Fig. 3C-D**), suggesting that RA was not sufficient to convert L cones to M cone fate. For most experimental conditions, we observed minimal differences between densities of M and L cones (**Fig. S2D**), suggesting that the changes in M to L cone ratios were primarily due to changes in cell fate specification, rather than dramatic loss or gain of cone subtypes. As the ‘Late RA’ condition did not affect M vs. L cone subtype specification (**Fig. 3C, 3G**), but did increase cone density observed on day 200 (**Fig. S2D**), addition of RA late in organoid development likely improves cone survival but not fate specification. Together, these data suggest that RA is sufficient to induce M cones and suppress L cones early in retinal organoid development, but not at later time points when cones are already specified.

### Natural variation in cone ratios is associated with RA signaling regulation

In parallel, we evaluated natural variation in the ratios of M and L cones in a human population sample. M and L cone ratios vary widely among people with normal color vision (*11*, *23*–*26*). We quantified cone ratios using flicker-photometric electroretinogram (FP-ERG), which measures and analyzes spectral sensitivities to 35 degrees of eccentricity (*23*), in 738 males with normal color vision. We studied males because variation at the X-linked *L/M-opsin* gene locus is a primary site for mutations causing red-green color blindness, and an increase in variation could be revealed in hemizygous conditions (*27*–*29*). We observed a range of L cone ratios (mean=59.1%; s.d.=16.9%; **Fig. 3H**). This extensive variation in the L and M cone ratios is in sharp contrast with the modest variation observed in the S cone ratio (8-12%) (*30*–*36*).

To identify a potential genetic basis of this variation in L and M cone ratios, we used a targeted sequencing approach to test for associations between cone cell ratios and genetic variation in mechanistic regulators of cone specification. Our sequencing targeted 21 gene regions, including the *L/M-opsin* gene locus (*OPN1LW/MW*), the *S*-*opsin* gene (*OPN1SW*), 3 RA regulatory genes (*RARA*, *NR2F2/COUP-TFII*, *CYP26A1/C1*), and 16 other genes with putative photoreceptor specification roles (*DIO1*, *DIO2*, *DIO3*, *PIAS3*, *RXRG*, *RORB*, *NR2F1/COUP-TFI*, *OTX1*, *NEUROD1*, *THRB*, *NRL*, *OTX2*, *ONECUT1*, *NR2E3*, *ONECUT2*, *SALL3*) (**Fig. S3**). We generated custom-designed oligos and used a solution-based capture method for target enrichment. We barcoded and pooled samples, which we sequenced with 125 bp paired-end reads. We mapped the reads, genotyped the samples, and evaluated associations with L cone ratios (**Fig. 3K**).

After alignment and genotyping, we scanned for association between variants in these genes and L cone ratios (**Fig. 3K**). We identified a significant association between SNP rs372754794 (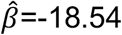, p-value=1.67 × 10^-5^) and variation in the L cone ratio (**Fig. 3I-K**). This SNP, which occurs within the targeted sequence spanning *NR2F2/COUP-TFII*, lies in an intron in a non-coding RNA (*NR2F2-AS1*) immediately upstream of *NR2F2* (**Fig. S4A-B**). The minor (G) allele, which is associated with reduced L cone ratio, was nearly exclusive to African American and African subjects within our sample (MAF=0.0359; **Fig. S4C**) and exhibited similar patterns of frequency differentiation in external data from the 1000 Genomes Project (*37*). The association with L cone ratio was robust even when restricting analysis to African American subjects (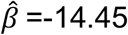, p-value=1.44 × 10^-4^) (**Fig. S4D**). Within African American populations from the 1000 Genomes Project, rs372754794 exhibited only modest linkage disequilibrium (LD) with other nearby variants (**Fig. S4B**), such that the associated haplotype is confined to the *NR2F2-AS1* transcript.

NR2F2 is a nuclear receptor that mediates RA signaling (*38*), which is expressed in the human retina and retinal organoids during development (**Fig. S5A-B**) (*16*). The associated SNP lies in a region of high transcription factor binding identified as a putative enhancer that is predicted to interact and regulate the promoters of *NR2F2* and/or *NR2F2-AS1* (**Fig. S5C-D).** Alternatively, as part of a lncRNA transcript, the SNP could affect *NR2F2-AS1-*mediated regulation of *NR2F2* or other genes. The observed association at NR2F2 is consistent with a role for RA signaling in M vs. L cone subtype specification.

The second strongest associated SNP (rs36102671) approached but did not achieve statistical significance upon multiple testing correction (Bonferroni correction p-value threshold=1.73 × 10^-5^). This SNP lies within the first intron of the *RARA* gene, which encodes a receptor for RA (**Fig. 3K**). The minor (A) allele, which segregates at low allele frequencies in globally diverse populations (**Fig. S6A**), is associated with decreased L cone ratios (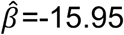, p-value=6.54 × 10^-5^). Within European populations, LD at this locus extends over a ~200 Kb region of chromosome 17, including *RARA* and 7 other genes (*R*^2^>0.7). The SNP exhibits an association with altered splicing of RARA based on RNA sequencing of whole blood (**Fig. S6B**) (*39*), suggesting that the haplotype modulates RARA function.

Together, the association of RA-related candidates with changes in M and L cone ratios is consistent with a role for RA signaling in the specification of M and L cone subtypes in humans.

## Discussion

We found that *M*-*opsin*, but not *L*-*opsin*, is expressed in early developing fetal retinas. In adults, the earlier-born central and middle regions contain a mix of M and L cones, whereas the later-born periphery is enriched for L cones. RA-synthesizing *ALDHs* are highly expressed early in retinal development and decrease over time, suggesting that RA signaling is high early. RA is sufficient to promote M cone fate and suppress L cone fate in immature cones. Together, we propose that RA signaling is high early in human retinal development to promote the generation of M cones and low later to yield the generation of L cones (**Fig. 3L**).

Our observations suggest that RA signaling influences cone fate in immature cones. In human fetal retinas, *M*-*opsin* mRNA expression is readily observed on day 115, whereas expression of RA synthesis genes (i.e., *ALDHs*) is highest on day ~55 and then decreases. In human retinal organoids, addition of RA before day 60, well before opsin mRNA expression, induces M cones and suppresses L cones. RA signaling may act at the gene locus to regulate opsin expression directly or function upstream to influence M and L cone fates.

As (1) *M*-*opsin* or *L*-*opsin* were exclusively expressed in cones in fetal and adult retinas and (2) RA promoted M cone fate early but did not convert L cones into M cones late, M cones and L cones likely arise directly from immature cones and do not undergo hybrid states. An alternative is that all cones initially express *M*-*opsin* and then a subset switch and express *L*-*opsin*, perhaps undergoing a period of co-expression during late fetal development, a timepoint which was experimentally inaccessible. Though these details require further interrogation, our data clearly support temporal generation and regulation of M and L cone fates in the human retina.

The arrangement of the LCR regulatory element and the *L-* and *M*-*opsin* genes in humans resembles an independently evolved gene array in zebrafish (*40*). In zebrafish, long wavelength opsin expression is controlled by RA, which promotes expression of the proximal opsin gene late in development (*41*). This contrasts with our findings that RA signaling promotes expression of the distal opsin early in humans. These observations suggest differences in the evolution of the regulation of RA signaling and the *cis*-regulatory logic controlling opsin expression, consistent with the independent origin of these opsin gene clusters.

Studying human development poses specific challenges including limited access to tissue, a lack of a controlled genetic background, and an inability to experimentally manipulate developing tissue. Vertebrate model organisms provided numerous insights into cell fate specification during retinal development. However, differences between these species and humans prevented interrogation of questions that can only be addressed in developing human tissue. Our studies show that human retinal organoids are a powerful model system to investigate developmental processes that are unique to humans and Old World monkeys.

## Supplemental Figure Legends

**Supplemental Fig. 1.**
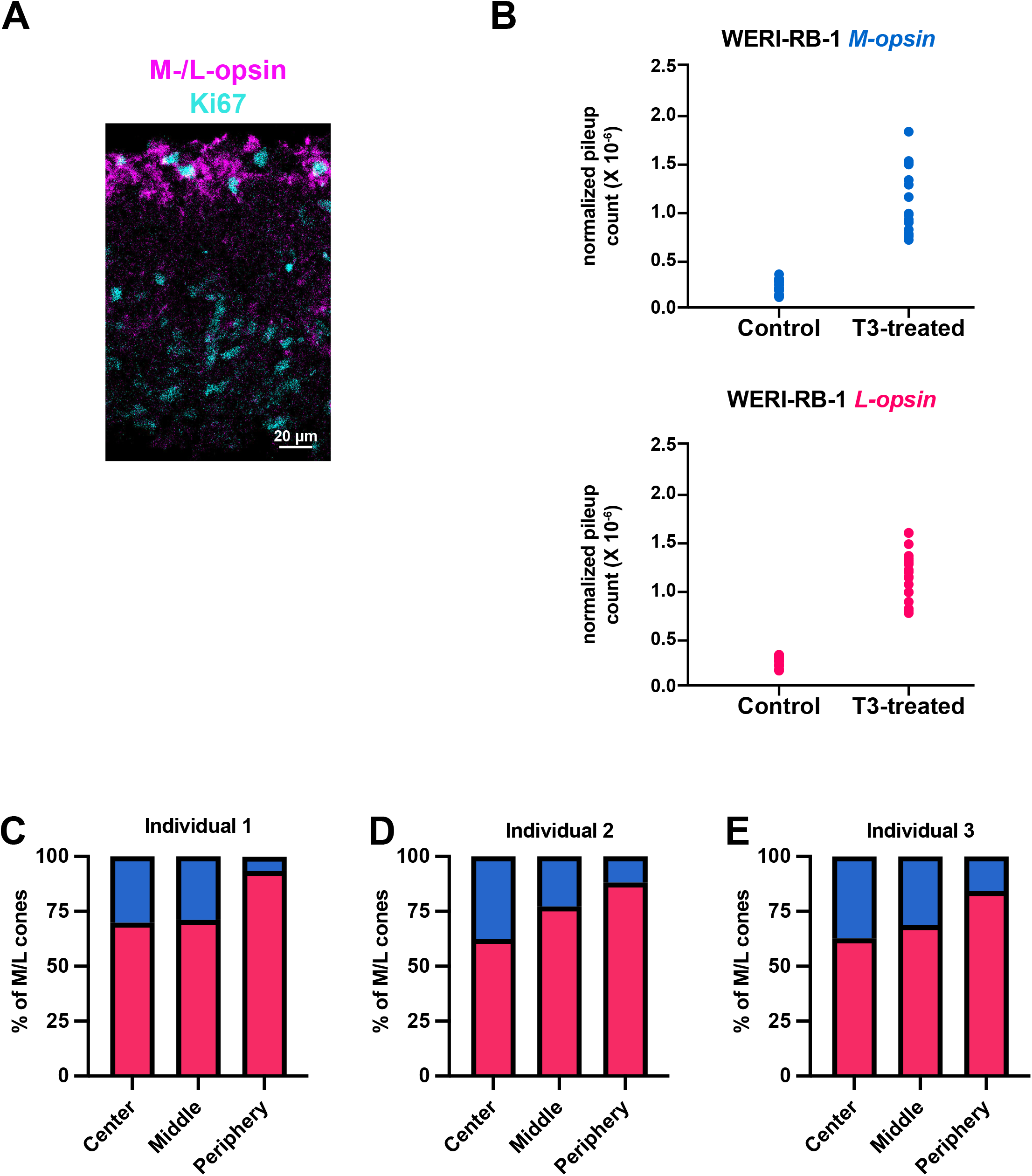
M-opsin and L-opsin expression in fetal and adult retinas. **(A)** 20 μm section of a fetal day 122 retina with expression of M-/L-opsin protein (magenta) and Ki67 (cyan). M-/L-opsin protein is observed in the outer nuclear layer. Ki67 expression indicates proliferating cells. **(B)** *M*- and *L*-*opsin* expression in control- and T3-treated Weri-Rb-1 retinoblastoma cells (N=1 experiment). The Weri-Rb-1 retinoblastoma cell line expresses *M*- and *L*-*opsin* at low levels (*42*). T3, the active form of thyroid hormone, induces *M*- and *L*-*opsin* expression in Weri-Rb-1 cells (*15*, *43*). Values indicate total pileup count normalized to total read count. Each data point indicates one nucleotide difference in *M*- or *L*-*opsin*. **(C-E)** Ratios of M and L cones as percent of M/L total cones for each individual. n>850 cones for each region for each individual. Averages ratios in (**Fig. 2J**).

**Supplemental Fig. 2.**
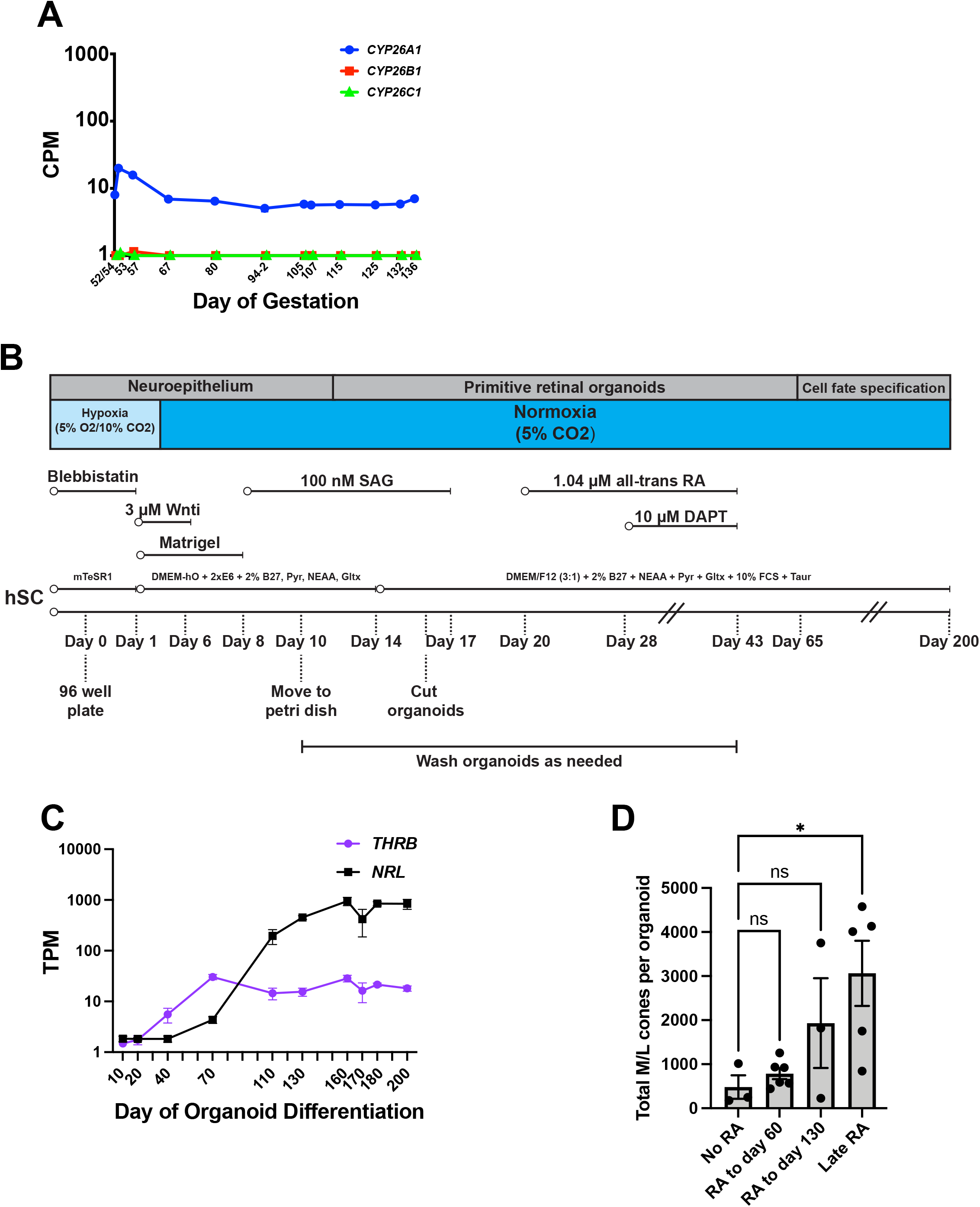
Organoid differentiation experiments and *CYP26A1/B1/C1* expression during fetal retinal development. **(A)** Expression of *CYP26A1, CYP26B1*, and *CYP26C1* in fetal human retinas. CPM=log counts per million. Analyzed from (*16*). **(B)** Human retinal organoid protocol. Standard conditions for human retinal organoid differentiation, adapted from (*15*). **(C)** Expression of *THRB* (cone marker) and *NRL* (rod marker) during retinal organoid development. TPM=transcripts per million. Analyzed from (*15*). **(D)** No significant differences in overall densities of M + L cones at day 200 in early RA treatment conditions (as in **Fig. 3C-F**) (Dunnett’s multiple comparison’s test, day 60 p=0.98, day 80 p=0.40, day 130 p=0.32). Significant difference in ‘No RA’ and ‘Late RA’ conditions (Dunnett’s multiple comparison’s test p-value=0.0182). Error bars indicate SEM. Individual circles represent individual organoids. One-way ANOVA with Dunnett’s multiple comparisons test ‘No RA’ versus ‘Late RA’ p<0.05.

**Supplemental Fig. 3.**
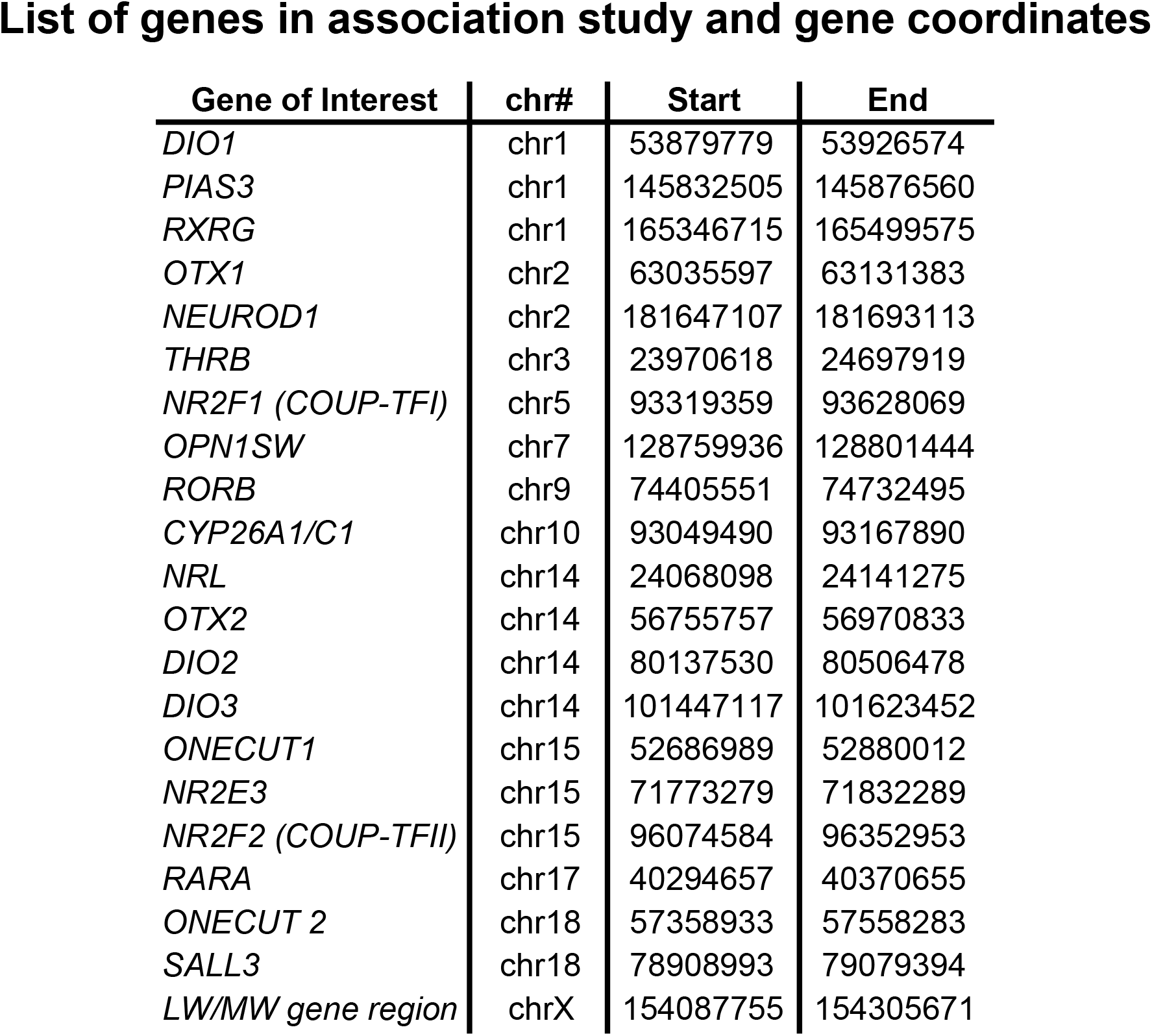
Genes tested in association study. Gene names and corresponding chromosome coordinates for assembly hg38.

**Supplemental Fig. 4.**
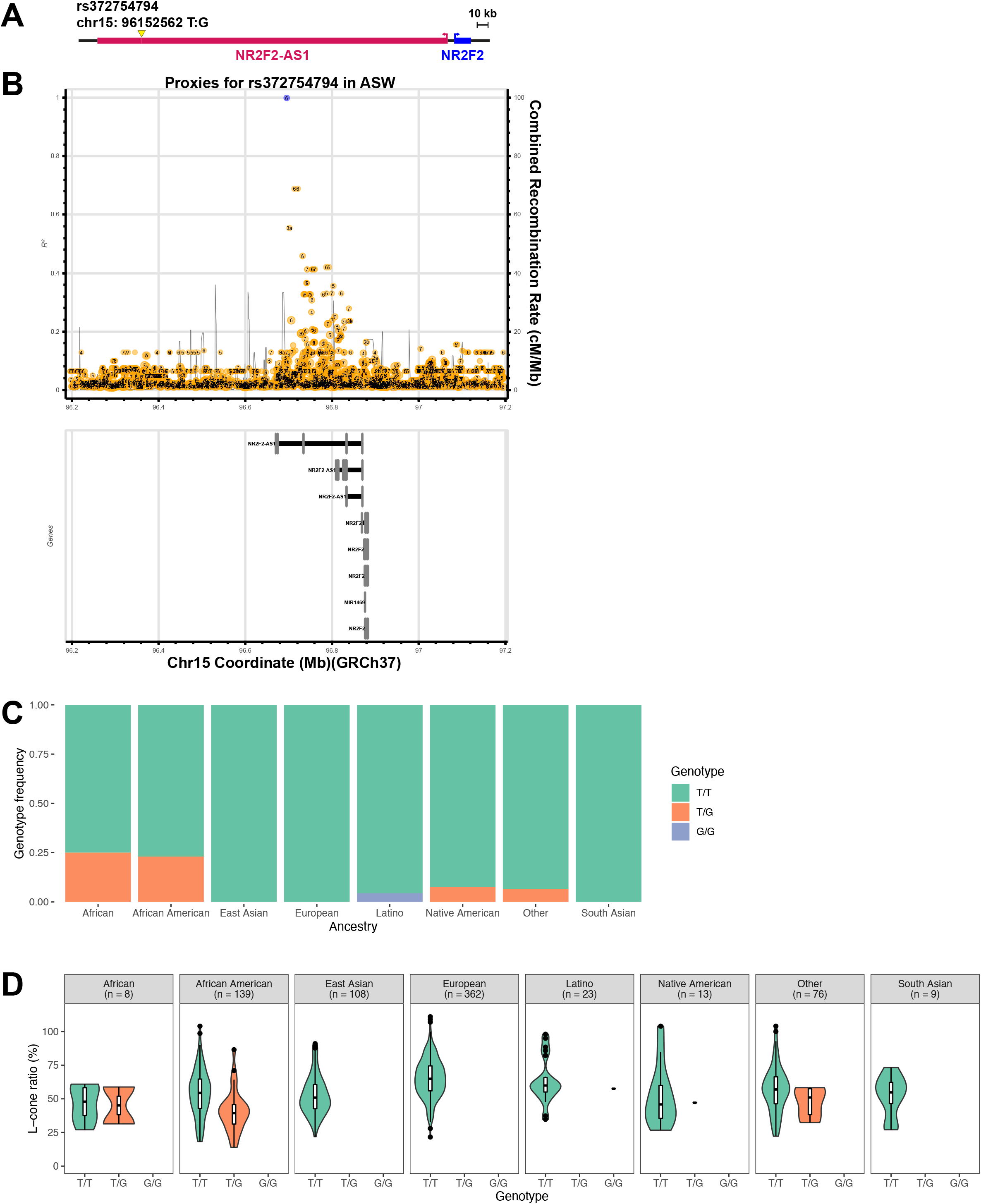
The rs372754794 SNP at the *NR2F2/NR2F2-AS1* locus is associated with differences in cone ratio. **(A)** *NR2F2* gene locus. *NR2F2* gene (pink); *NR2F2* antisense RNA (blue). Yellow arrow denotes the location of the SNP. **(B)** Local LD plot for rs37275494 based on data from an African American (ASW) population from the 1000 Genomes Project. Top indicates variants. Bottom indicates gene predictions. **(C)** Minor (G) allele ratio by ancestry. **(D)** Association of L:M ratio and rs372754794 stratified by self-reported ancestry.

**Supplemental Fig 5.**
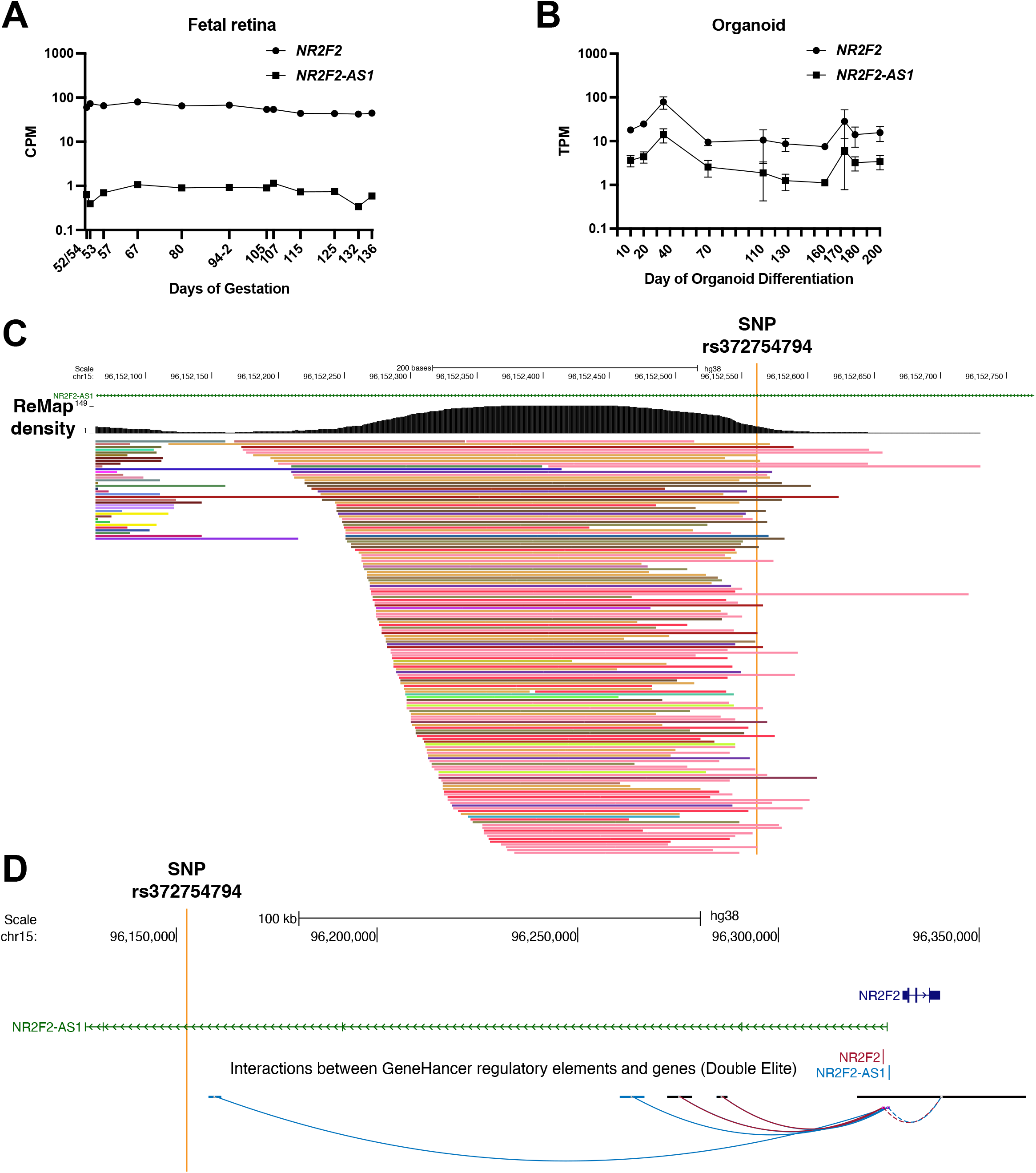
Expression of *NR2F2* and *NR2F2-AS1* during development; The rs372754794 SNP at the *NR2F2/NR2F2-AS1* locus lies in a putative regulatory region. **(A)** Expression of *NR2F2* and *NR2F2-AS1* in human fetal retinas, analyzed from (*16*). **(B)** Expression of *NR2F2* and *NR2F2-AS1* in human retinal organoids, analyzed from (*15*). **(C)** ReMap ChIP-seq database shows that the rs372754794 SNP lies in an enhancer based on transcription factor binding. Each colored line indicates ChIP-seq binding data for a different transcriptional regulator. The ReMap density show the density of the peaks overlap. **(D)** GeneHancer database shows that the rs372754794 SNP neighbors a region predicted to physically interact and regulate *NR2F2* and/or *NR2F2-AS1*.

**Supplemental Fig 6.**
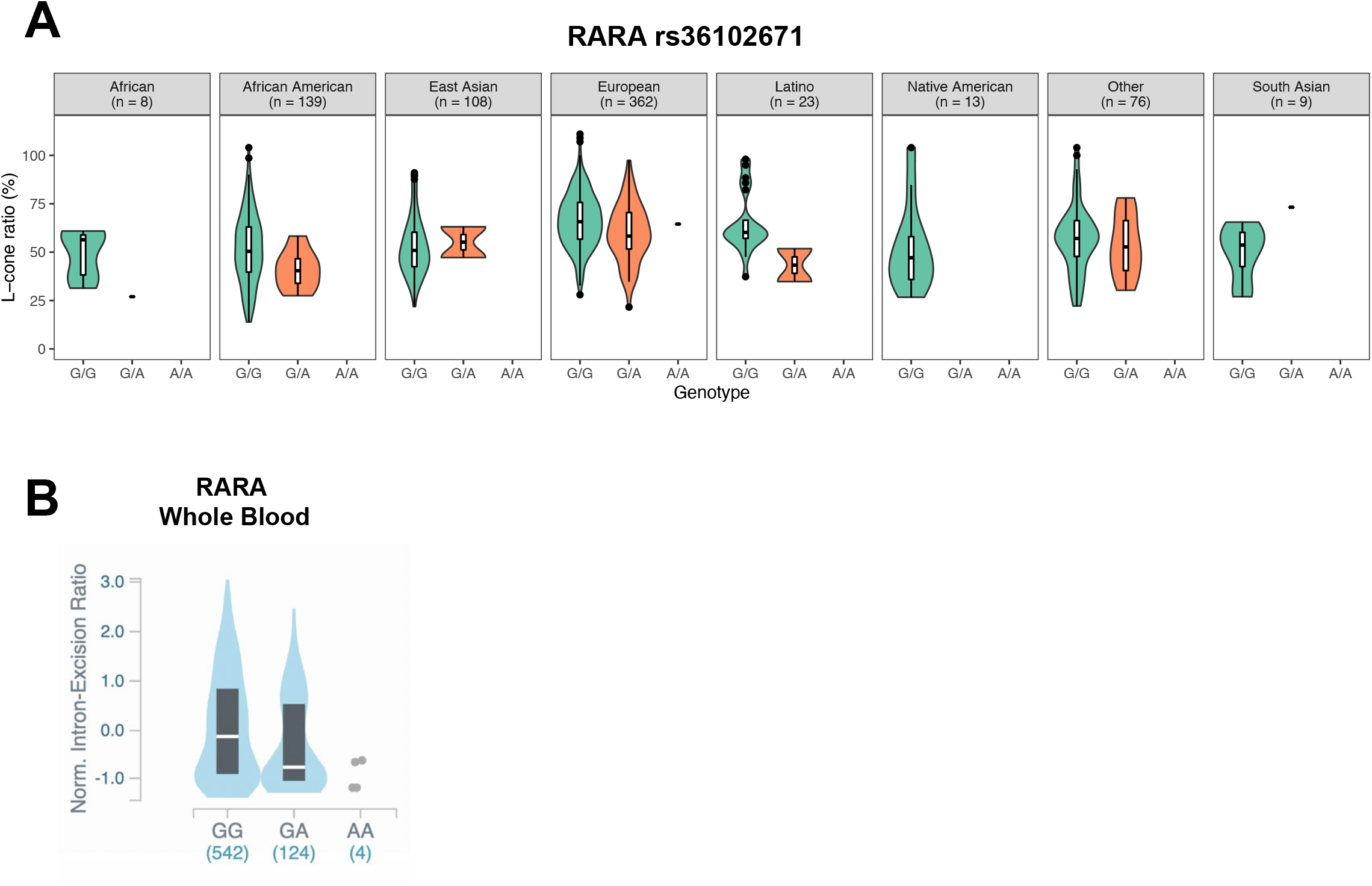
The rs36102671 SNP at the *RARA* locus is associated with differences in cone ratio. **(A)** Association of L:M ratio and rs36102671 stratified by self-reported ancestry. **(B)** rs36102671 association with altered splicing of RARA in whole blood from the Genotype Tissue Expression Project (GTEx).

## Supplemental Information

### Materials and Methods

#### Cell Lines

Cell lines have not been authenticated

#### HEK293 cell line maintenance

HEK293 cells were maintained in DMEM 4.5 g/L D-glucose, L-glutamine (11965084, Gibco) + 10% Fetal Bovine Serum (16140071, Gibco) + 1X Penicillin-Streptomycin (30-002-CI, Corning) at 37°C in a HERAcell 150i or 160i 5% CO_2_ incubator (Thermo Fisher Scientific). Cells were split 1:4 – 1:10 every 2-3 days when approximately 80% confluent and kept in 6-well dishes or uncoated flasks using Trypsin (0.25%)-EDTA (0.02%) in HBSS (118-093-721, Quality Biological).

#### Transfection of opsin cDNA into HEK293 cells

HEK293 cells were passaged 24 hours prior to be <60% confluent on the day of transfection. Transfection was done with the Lipofectamine^®^ 3000 transfection kit (L3000001, Invitrogen). For a 10 cm dish, 7.5 μL lipofectamine 3000 was mixed with 125 μL Opti-MEM (31985062, Thermo Fisher Scientific) at room temperature for 5 minutes. 5 μL of P3000, 0.5-1 μg DNA (**Table 2**), and 125 μL Opti-MEM was mixed at room temperature for 5 minutes as per manufacturer instruction. The two solutions were mixed and incubated at room temperature for 10 minutes. Media was aspirated from HEK293 cells and replaced with 8 mL of pre-warmed Opti-MEM. Lipofectamine mixture was added to cells and incubated at 37°C in a HERAcell 150i or 160i 5% CO_2_ incubator for 4-6 hours. Transfection media was aspirated and replaced with HEK293 maintenance media (DMEM/10% FBS/1X Penn/Strep) and cells were allowed to recover overnight. Cells were adhered to slides coated with 10% Poly-L-lysine solution (0.1% w/v in H_2_O) (P8920, Sigma) for 1-3h at 37°C in a HERAcell 150i or 160i 5% CO_2_ incubator (Thermo Fisher Scientific). Cells were washed in 1x PBS and fixed in 10% neutral buffered formalin (HT501128, Sigma) for 30 minutes. Slides were either used immediately or ethanol dehydrated and stored at −80°C.

**Table 1:** Genotypes of cell lines.

| Cell line | Sex | Source | Figures |
| --- | --- | --- | --- |
| HEK293 | Female | Kuruvilla lab, Johns Hopkins | **1C-J** |
| Weri-Rb-1 | Female | ATCC | **S1B** |
| H7 ESC | Female | WA07, WiCell | **3C-G, S2D** |
| H7 iCas9 ESC | Female | Zack lab, Johns Hopkins | **3C-G, S2D** |
| EP1.1 iPSC | Female | Zack lab, Johns Hopkins | **S2C, S5B** |

Cell lines have not been authenticated.

**Table 2:** Opsin cDNA plasmids.

| Plasmid | Source | Figure |
| --- | --- | --- |
| OPN1MW NM_000513.2 | GenScript | **1C-J** |
| OPN1LW NM_020061.5 | GenScript | **1C-J** |

#### BaseScope RNA in situ hybridization

BaseScope RNA *in situ* hybridization was performed according to manufacturer’s instructions (Advanced Cell Diagnostics) and modified with several changes. The entire protocol is detailed below. Sections of human, fetal, or organoid tissue were allowed to come to room temperature from storage at −80C and rehydrated in 1X PBS. Samples were pretreated according to manufacturer’s instructions: 10 minutes in RNAscope Hydrogen Peroxide followed by a wash in dH_2_O and then 2 washes in 1X PBS. The following step is an RNAscope Protease step which varies depending on tissue type: for HEK293 cells RNAscope Protease III was applied at a 1:15 dilution in 1X PBS for 15 minutes in a humid chamber. RNAscope Protease IV was applied for 20 minutes in a humid chamber for organoids and human eye samples. Samples were then washed in 1X PBS twice. Probes were added (**Table 3**) at the manufacturer suggested concentration to samples in the HybEZ Humidity Control rack with lid and insert into the HybEZ oven for 2 hours at 40°C. Samples were washed twice in 1x RNAscope wash buffer for 2 min. Amplification and development washes were done with the manufacturer reagents without change to the recommended concentrations. All washes were done in the HybEZ Humidity Control rack with lid and insert, either at room temperature (RT) or at 40°C in the HybEZ oven at the temperatures and time lengths according to **Table 4**, with 2X 2 min washes in 1X RNAScope Buffer conducted between each reagent wash.

**Table 3:** BaseScope *in situ* probes.

| Reagent type | Reagent | Source | Identifier | Additional information |
| --- | --- | --- | --- | --- |
| BaseScope probe | BA-Hs-OPN1MW-1zz-st1 | Advanced Cell Diagnostics | 714201 | bp 902-941 NM_000513.2 |
| BaseScope probe | BA-Hs-OPN1LW-1zz-st-C2- | Advanced Cell Diagnostics | 716081-C2 | bp 880-921 NM_020061.5 |

**Table 4:** BaseScope assay reagent washes.

| Reagent | Temperature (°C) | Time (min) |
| --- | --- | --- |
| AMP 1 | 40 | 30 |
| AMP 2 | 40 | 30 |
| AMP 3 | 40 | 15 |
| AMP 4 | 40 | 30 |
| AMP 5 | 40 | 30 |
| AMP 6 | 40 | 15 |
| AMP 7 | RT | 30 |
| AMP 8 | RT | 15 |
| RED-A/B | RT | 10 |
| AMP 9 | 40 | 15 |
| AMP 10 | 40 | 15 |
| AMP 11 | RT | 30 |
| AMP 12 | RT | 15 |
| GREEN-A/B | RT | 10 |

Slides were baked for a minimum of 30 minutes at 65°C in the HybEZ oven or until dry. Samples were preserved with VectaMount (Vector Laboratories, H-5000) mounting media and a sealed 1.5 coverslip.

#### Imaging HEK293 RNA in situ hybridization

HEK293 cells were imaged using an EVOS XL Core Cell imaging system on brightfield with a 20x air objective (ThermoFisher).

#### M/L-opsin immunohistochemistry and in situ hybridization BaseScope on HEK293 cells and adult human retina sections

*M*- and *L*-*opsin* transfected HEK293 cells were allowed to come to room temperature after storage in 100% EtOH at 4°C. Adult sections were allowed to come to room temperature from storage at −80C and dehydrated in a stepwise ethanol dehydration from 50%, 70% and 100% EtOH. Samples were pretreated according to ACD Biotechne manufacturer’s instructions: 10 minutes of RNAscope^®^ Hydrogen Peroxide at room temperature, followed by a wash in dH2O and then 2 washes in 1X PBS. Slides were then acclimatized for 10 seconds in 99°C dH_2_0, and then transferred to 99°C RNAscope^®^ 1X Target Retrieval Reagent for 15 minutes using the Oster Double Tiered Food Steamer Warmer 5712. Slides were then transferred into dH_2_0 for 15 seconds, and then transferred to 0.1% TWEEN (Sigma P7949). A primary antibody targeting both OPN1MW and OPN1LW (antirabbit Millipore ab5405 #3823289) was diluted 1:200 in Co-detection Antibody Diluent and added to the slides. All slides were incubated overnight in a humidity-controlled tray at 4°C. Primary antibodies were washed off with two washes of 0.1% TWEEN for 2 minutes, and then post-fixed by submerging in 10% Neutral Buffered Formalin for 30 minutes at room temperature. Fixative was removed by four washes of 0.1% TWEEN for 2 minutes each. For HEK293 cells, RNAscope^®^ Protease III was applied at a 1:15 dilution in 1X PBS for 15 minutes in a humid chamber. For retina sections, RNAscope^®^ Protease IV was applied for 20 minutes in a humid chamber. Samples were washed in 1X PBS twice. Slides were washed twice in RNAscope^®^ 1X wash buffer for 2 minutes each. CoDetection Blocker was applied to the slides and incubated for 15 minutes at 40°C. Slides were then washed twice in RNAscope^®^ wash buffer for 2 minutes each, and then washed in 0.1% TWEEN for 2 minutes. Donkey anti-rabbit 647 secondary antibody (Invitrogen Thermo Fisher Scientific A31573 #2181018) was diluted 1:400 in Co-detection Antibody Diluent and applied to the samples overnight in a humidity control tray at 4°C. Slides were then washed twice in 0.1% TWEEN for 2 minutes each, and co-stained in hoechst 33342 (Biotium 40046 #10H0212, diluted 1:2000 in 1X PBS).

#### Imaging of M/L-opsin IHC and ISH

Slides were mounted in 1X PBS and the IHC staining was imaged the next day using either a Zeiss LSM 980 inverted microscope with an Axiocam 506 color camera system and a C Apo 40x/1.2 W DICIII lens or an Olympus IX81 inverted microscope equipped with a Hamamatsu Orca 4.0 v3 monochrome camera.

#### ISH BaseScope of M and L-opsin in human adult retina and HEK293 cells

Coverslips from previous IHC assay were removed by soaking for 10 minutes in 5X SSC (Invitrogen by Thermo Fisher Scientific AM9763). BaseScope probes were then applied and amplified using the aforementioned protocol (“<u>BaseScope RNA in situ hybridization”)</u>. Nuclei were stained using hoescht 33342 (Biotium 40046 #10H0212, diluted 1:2000 in 1X PBS).

#### Imaging of ISH BaseScope M- and L-opsin in human adult retina and HEK293 cells

ISH BaseScope colorimetric staining was then imaged the day after protocol completeion in brightfield using either a Zeiss LSM 980 inverted microscope with an Axiocam 506 color camera system and a C Apo 40x/1.2 W DICIII lens, or an Olympus IX81 inverted microscope equipped with a Hamamatsu Orca 4.0 v3 monochrome camera. A Cambridge Research & Instrumentation, Inc. (CRi) Micro*Color tunable RGB filter was inserted in the light path to allow fast sequential acquisition of red, green, and blue images, which were merged to form the final RGB dataset.

#### Human adult retina tissue

Donor samples were acquired from the National Disease Research Interchange (NDRI) between 12.5-15.5 hours postmortem and were flash frozen on dry ice after enucleation and stored at −80°C. A human eye was allowed to come to room temperature in 1X PBS. The anterior portion of the eye (cornea, iris, lens) was removed. The posterior pole of the eye was butterflied in a petri dish, with 4-5 cuts made from the anterior ciliary body to midway to the posterior, such that the retina and musculature laid flat on the dish. A 5 mm punch biopsy tool (RBP-50, Robbins Instruments) was placed over the macula based on the presence of macular pigment in combination with presence of an avascular region to remove the central retina. Punches were taken immediately adjacent for the middle retina, and immediately adjacent for the periphery (**Fig. 2F**). Retinal punches were fixed for 45 minutes in 10% neutral buffered formalin (HT501128, Sigma) and washed in 1X PBS. The subsections of retina were mounted in Tissue-Tek O.C.T. compound (4583, Sakura), placed on dry ice to freeze, and stored at −80°C. The retina was cryosectioned in 20 μm sections. Slides were air dried for 6 hours to overnight followed by a post-fixation step of 15 minutes in 10% neutral buffered formalin (HT501128, Sigma) and washed in 1X PBS after drying. Slides were dried and stored at −80°C for no more than 3 months before use.

#### Imaging human adult RNA in situ hybridization

Samples were imaged in brightfield using a Zeiss LSM 980 inverted microscope with an Axiocam 506 color camera system using a C Apo 40x/1.2 W DICIII lens or an EVOS XL Core Cell imaging system on brightfield with a 20x air objective (ThermoFisher).

#### Human fetal retina tissue

Day 122 fetal retina (sex unknown) was enucleated and flash frozen on dry ice and stored at −80°C as a gift from the Reh lab at the University of Washington. The fetal eye was allowed to come to room temperature in 1X PBS. The anterior portion of the eye (lens, cornea, iris) were removed with scissors. The posterior portion of the eye was fixed overnight in 10% neutral buffered formalin (HT501128, Sigma) and washed in 1X PBS. The whole posterior eye was mounted in Tissue-Tek O.C.T. compound (4583, Sakura), placed on dry ice to freeze, and stored at −80°C. The eye was cryosectioned in 20 μm sections starting from the anterior side to the posterior, such that each section contains the nasal/temporal and dorsal/ventral information at each position along the anterior/posterior axis. Sections were fixed in a post-fixed for 15 minutes in 10% neutral buffered formalin (HT501128, Sigma) and washed in 1X PBS, then air dried overnight. Slides were stored at −80°C and used within six months.

#### Imaging human fetal RNA in situ hybridization

Samples were imaged in brightfield using a Zeiss LSM 980 inverted microscope with an Axiocam 506 color camera system using a C Apo 40x/1.2 W DICIII lens or an EVOS XL Core Cell imaging system on brightfield with a 20x air objective (ThermoFisher).

#### IHC of fetal human retina sections

Day 122 human fetal retina sections were stained using the above IHC staining protocol (“M/L-opsin IHC and ISH BaseScope on HEK293 cells and adult human retina sections”) with the following adjustments: In addition to OPN1MW and OPN1LW, primary antibodies targeting Ki67 (anti-mouse Santa Cruz Biotechnology sc-23900 #A1520) and OPN1SW (anti-chick, gifted by J. Nathans), diluted 1:200 in Co-detection Antibody Diluent) were applied to the fetal human retina sections. The following secondaries were used: donkey anti-rabbit 647 (Invitrogen Thermo Fisher Scientific A31573 #2181018), donkey antimouse 555 secondary antibody (Invitrogen Thermo Fisher Scientific A31570 #204536) and donkey anti-chick 488 secondary antibody (703-545-155 #156558 Jackson ImmunoResearch Laboratories Inc) was diluted 1:400 in Co-detection Antibody Diluent. Samples were imaged using a laser scanning confocal Zeiss LSM 980 inverted microscope using a C Apo 40x/1.2 W DICIII lens.

#### Weri-Rb-1 cell line maintenance

Weri-Rb-1 retinoblastoma cells were maintained in RPMI 1640 Medium (11875135, Gibco) + 10% Fetal Bovine Serum (16140071, Gibco) + 1X Penicillin-Streptomycin (30-002-CI, Corning) at 37°C in a HERAcell 150i or 160i 5% CO_2_ incubator (Thermo Fisher Scientific). Cells were passaged every 4 days at ~1 x 10^5^ – 2 x 10^6^ cells/mL in uncoated flasks by pelleting at 150g for 5 minutes and resuspending in fresh media.

#### Bulk RNA sequencing samples

Fetal retina bulk RNA sequencing data were processed in Hoshino *et al*. 2017 [1]. Adult retina sequencing data were processed in Pinelli *et al*. 2016 [2]. Weri-Rb-1 samples were grown in control media or T3-treated media (100 nM T3 (T6397, Sigma) in RPMI + supplement media) for four days (N=1). EP1 iPSC-derived organoids were previously grown and analyzed for Eldred *et al*. 2018 at time points ranging from day 10 to day 250 of differentiation [3]. Samples were taken at day 10 (N=3), day 20 (N=2), day 35 (N=3), day 69 (N=3), day 111 (N=3), day 128 (N=3), day 158 (N=2), day 173 (N=3), day 181 (N=3), day 200 (N=3), and day 250 (N=3). RNA from individual samples was extracted using the Zymo Direct-zol RNA Microprep Kit (R2062, Zymo Research) according to manufacturer’s instructions. Libraries were prepared using the Illumina TruSeq stranded mRNA kit and sequenced on an Illumina NextSeq 500 with single 75 bp reads.

#### Bulk RNA sequencing analysis

Quantification of opsin expression in human fetal retina samples from [1, 2] using code deposited https://github.com/bbrener1/johnston_retina: Expression of opsin genes was directly quantified from FASTQ files using Kallisto 0.44.0. The kallisto index was generated from the Gencode V27 human transcriptome, with the default kmer setting of 31. Reads were quantified in single-ended mode with a specified length of 75 and standard deviation of 10. Quantification of opsin expression in organoid data: Organoid data was previously quantified and described in (1). Expression levels were quantified using Kallisto (version 0.34.1) with the following parameters: “-b 100 -l 200 -s 10 -t 20 --single”. The Gencode release 28 comprehensive annotation was used as the reference transcriptome [4]. For the analysis of opsin pileups, to examine the reads mapped to each opsin directly, reads were aligned to the opsin transcripts using Bowtie 2.3.4.3, and aligned files were processed with samtools 1.7 to produce pileups around selected locations in the transcripts. Pileups are a text-based format that quantifies base calls of aligned reads to a reference genome. For the quantification of human samples, a Bowtie2 index was generated from sequences ENST00000595290.5 and ENST00000369951.8 using the default settings. Pileups were generated for transcript positions given in **Table 5**.

**Table 5:** Transcript positions for pileup analyses.

| Exon | Base |  | Transcript position |  | Exon Position | Absolute Position |  |
| --- | --- | --- | --- | --- | --- | --- | --- |
|  | LW | MW | LW | MW |  | LW | MW |
| 2 | C | T | 254 | 276 | 81 | 154150737 | 154187851 |
|  | A | G | 360 | 382 | 187 | 154150843 | 154187957 |
|  | A | G | 391 | 413 | 218 | 154150874 | 154187988 |
|  | C | A | 407 | 429 | 234 | 154150890 | 154188004 |
| 4 | T | C | 749 | 771 | 110 | 154154684 | 154191798 |
|  | G | A | 757 | 779 | 118 | 154154692 | 154191806 |
|  | C | G | 758 | 780 | 119 | 154154693 | 154191807 |
|  | T | C | 759 | 781 | 120 | 154154694 | 154191808 |
|  | A | G | 766 | 788 | 127 | 154154701 | 154191815 |
| 5 | A | G | 880 | 902 | 75 | 154156369 | 154193483 |
|  | T | C | 883 | 905 | 78 | 154156372 | 154193486 |
|  | T | G | 885 | 907 | 80 | 154156374 | 154193488 |
|  | G | A | 888 | 910 | 83 | 154156377 | 154193491 |
|  | A | T | 890 | 912 | 85 | 154156379 | 154193493 |
|  | G | T | 895 | 917 | 90 | 154156384 | 154193498 |
|  | C | A | 901 | 931 | 104 | 154156398 | 154193512 |
|  | A | G | 905 | 935 | 108 | 154156402 | 154193516 |
|  | T | C | 940 | 970 | 143 | 154156437 | 154193551 |
|  | G | C | 944 | 974 | 147 | 154156441 | 154193555 |
|  | A | T | 978 | 1008 | 181 | 154156475 | 154193589 |

#### Stem cell line maintenance

H7 ESC (WA07, WiCell) and episomal-derived EP1.1 iPSC (Zack lab, Hopkins) were used for retinal organoid differentiation. Pluripotency of EP1.1 cells was evaluated previously with antibodies for NANOG, OCT4, SOX2, and SSEA2 [5]. Stem cells were maintained in mTeSR™1 (85857, Stem Cell Technologies) on 1% (v/v) Matrigel-GFR™ (354230, BD Biosciences) coated dishes and grown at 37°C in a HERAcell 150i or 160i 10% CO_2_ and 5% O_2_ incubator (Thermo Fisher Scientific). Cells were passaged every 4-5 days according to confluence as in *[5]*. Cells were passaged with Accutase (SCR005, Sigma) for 7-12 minutes to be dissociated to single cells. Cells in Accutase were added 1:2 to mTeSR™1 plus 5 μM Blebbistatin (Bleb, B0560, Sigma), pelleted at 150g for 5 minutes, and suspended in mTeSR™1 plus Bleb and plated at 5,000-15,000 cells per well in a 6-well plate. Media was replaced with mTeSR™1 48 hours following passage and every 24 hours until passaged again. To minimize cell stress, no antibiotics were used.

#### Organoid media

E6 supplement: 970 μg/mL Insulin (11376497001, Roche), 535 μg/mL holo-transferrin (T0665, Sigma), 3.20 mg/mL L-ascorbic acid (A8960, Sigma), 0.7 μg/mL sodium selenite (S5261, Sigma).

BE6.2 media for early retinal differentiation: 2.5% E6 supplement (above), 2% B27 Supplement (50X) minus Vitamin A (12587010, Gibco), 1% Glutamax (35050061, Gibco), 1% NEAA (11140050, Gibco), 1mM Pyruvate (11360070, Gibco), and 0.87 mg/mL NaCl in DMEM (11885084, Gibco).

LTR (Long-Term Retina) media: 25% F12 (11765062, Gibco) with 2% B27 Supplement (50X) (17504044, Gibco), 10% heat inactivated FBS (16140071, Gibco), 1mM Sodium Pyruvate,1% NEAA, 1% Glutamax and 1 mM taurine (T-8691, Sigma) in DMEM (11885084, Gibco).

#### Retinoic acid treatments

For organoids, 1.04 μM all-trans retinoic acid (ATRA; R2625; Sigma) was thawed fresh in LTR media every two days, protected from light, and never freeze-thawed.

#### Organoid differentiation and maintenance

Organoids were differentiated from H7 ESCs, H7iCas9 ESCs, or EP1.1 iPSCs (**Table** 6) as described in Eldred *et al*. 2018 with minor variations (**Fig. S3**) [3]. On day 0, pluripotent and well-maintained stem cells with minimal to no spontaneous differentiation were used for organoid aggregation. To aggregate, cells were passaged in Accutase (SCR005, Sigma) at 37°C for 12 min to ensure complete dissociation. Cells were seeded in 50 μLs of mTeSR1 (85857, Stem Cell Technologies) at 3,000 cells/well into 96-well ultra-low adhesion round bottom Lipidure coated plates (51011610, NOF) or ultra-low attachment microplate (7007, Corning). Corning plates were used upon the discontinuation of Lipidure plates. Cells were placed in hypoxic conditions (10% CO_2_ and 5% O_2_) for 24 hours to enhance survival. Cells naturally aggregated by gravity over 24 hours. On day 1, cells were moved to normoxic conditions (5% CO_2_). On days 1-3, 50 μLs of BE6.2 media containing 3 μM Wnt inhibitor (IWR1e: 681669, EMD Millipore) and 1% (v/v) Matrigel were added to each well. On days 4-9, 100 μLs of media were removed from each well, and 100 μLs of media were added. On days 4-5, BE6.2 media containing 3 μM Wnt inhibitor and 1% Matrigel was added. On days 6-7, BE6.2 media containing 1% Matrigel (354230, BD Biosciences) was added. On days 8-9, BE6.2 media containing 1% Matrigel and 100 nM Smoothened agonist (SAG: 566660, EMD Millipore) was added.

**Table 6:**
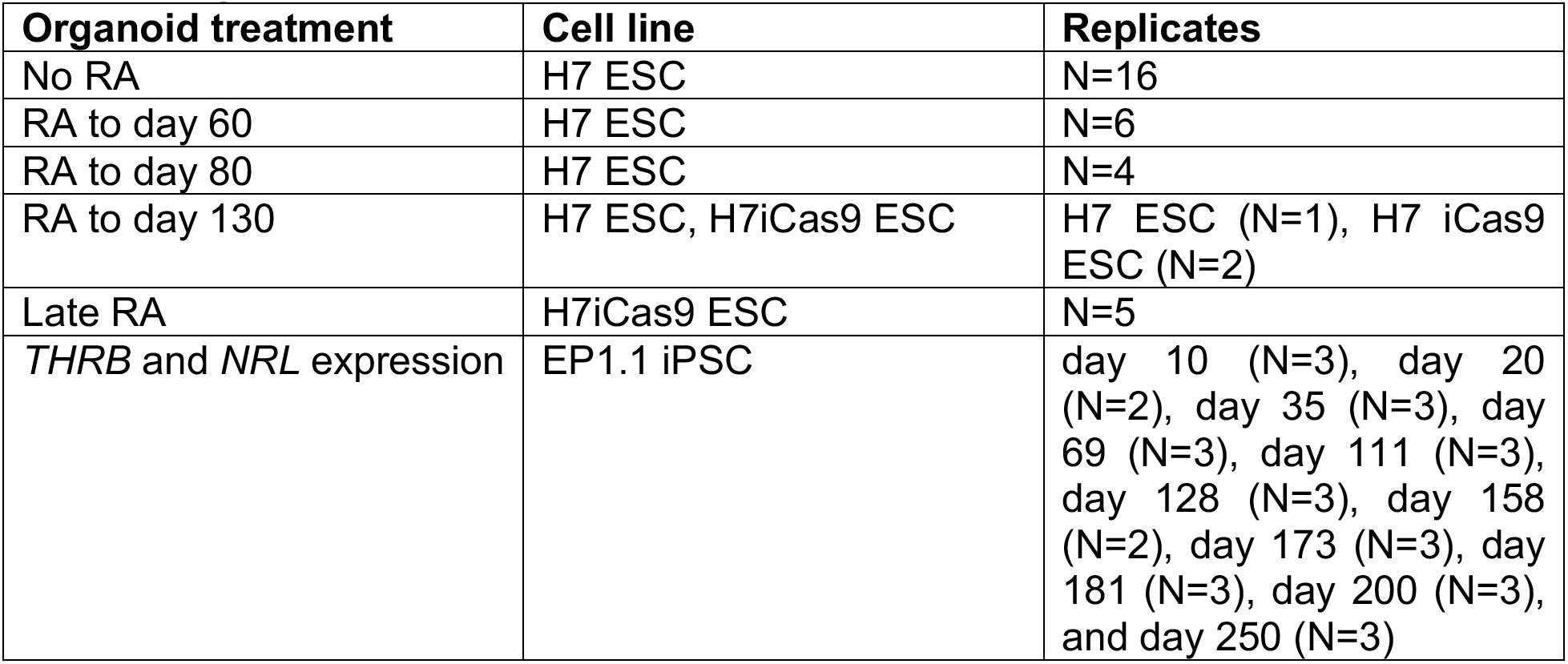
Organoid replicates and cell lines.

| Organoid treatment | Cell line | Replicates |
| --- | --- | --- |
| No RA | H7 ESC | N=16 |
| RA to day 60 | H7 ESC | N=6 |
| RA to day 80 | H7 ESC | N=4 |
| RA to day 130 | H7 ESC, H7iCas9 ESC | H7 ESC (N=1), H7 iCas9 ESC (N=2) |
| Late RA | H7iCas9 ESC | N=5 |
| *THRB* and *NRL* expression | EP1.1 iPSC | day 10 (N=3), day 20 (N=2), day 35 (N=3), day 69 (N=3), day 111 (N=3), day 128 (N=3), day 158 (N=2), day 173 (N=3), day 181 (N=3), day 200 (N=3), and day 250 (N=3) |

On day 10, aggregates were transferred to 15 mL tubes, rinsed 3X in 5 mL DMEM (11885084, Gibco), and resuspended in BE6.2 with 100 nM SAG in untreated 10 cm polystyrene petri dishes in a total of 12 mL of media. From this point on, media was changed every other day. Aggregates were monitored and manually separated if stuck together or to the bottom of the plate. On days 13-16, LTR media with 100 nM SAG was added. On day 16, retinal vesicles were manually dissected using sharpened tungsten needles, with cuts made to maximize the number of vesicles per organoid. On average, one to two cuts were made per organoid. After dissection, cells were transferred into 15 mL tubes and washed 3X with 5 mLs of DMEM (11885084, Gibco). On days 16-20, cells were maintained in LTR and washed 2X with 5 mLs of DMEM (11885084, Gibco), to remove dead cells, before being transferred to new plates. To increase survival and differentiation, 1.04 μM all-trans retinoic acid (ATRA; R2625; Sigma) was added to LTR medium from days 20-43. Additional time windows of 1.04 μM were added depending on experimental conditions. 10 μM gamma-secretase inhibitor (DAPT, 565770, EMD Millipore) was added to LTR from days 28-42. Organoids were grown at low density (10-20 per 10 cm dish) to reduce aggregation. Periodically, organoids were culled from the plate based on absence of clear laminal structure indicating proper retinal organoid growth.

#### Organoid preparation and cryosectioning

Organoids were fixed for 45 minutes in 10% neutral buffered formalin (HT501128, Sigma) and washed in 1X PBS. Organoids were placed in a 25% sucrose in 0.1 M phosphate buffer solution overnight, and then mounted in Tissue-Tek O.C.T. compound (4583, Sakura), placed on dry ice to freeze, and stored at −80C. Organoids were cryosectioned in 10 μm sections. Slides were air dried for 6 hours to overnight, then post-fixated for 15 minutes in 10% neutral buffered formalin (HT501128, Sigma) and washed in 1X PBS. Slides were dried and stored at −80°C and used within 3 months.

#### Imaging organoid RNA in situ hybridization

Samples were imaged in brightfield using a Zeiss LSM 980 inverted microscope with an Axiocam 506 color camera system using a C Apo 40x/1.2 W DICIII lens or an EVOS XL Core Cell imaging system on brightfield with a 20x air objective (ThermoFisher).

#### Association study subjects

All experiments involving human subjects were conformed to the principles expressed in the Declaration of Helsinki and were approved by the University of Washington Human Subjects Review Committee. Subjects were 738 males who had previously passed several standard tests for color vision deficiency, including Ishihara’s 24-plate test, Richmond HRR 2002 edition, and the Neitz Test of Color Vision. Subjects identified themselves as being either of African or African American ancestry (N=147), Asian ancestry (N=108), Caucasian ancestry (N=362), Latino ancestry (N=23), Mixed (N=62), Native American ancestry (N=13), South Asian ancestry (N=9), or Other/Not Specified (N=14).

#### Flicker photometric electroretinogram (FP-ERG)

The proportion of L cones, expressed as the percentage of L plus M cones that were L, were determined for each subject from a combination of genetic analysis and the flicker photometric electroretinogram (FP-ERG) as described in [6, 7]. Briefly, the spectral sensitivity of the individual was measured using control and test lights of specific wavelengths to illuminate a small portion of the retina. These conditions were designed to eliminate rod and S-cone contributions. While the test lights were flickered on and off, electrodes recorded the ERG neuronal responses to the stimulus. The spectral sensitivity was determined by adjusting the intensity of the test light until the ERG signal produced exactly matched that produced by the fixed intensity reference light. A correction factor was applied to the estimated proportion of L cones to account for the 1.5 times greater contribution to the FP-ERG signal for each M cone compared to each L cone [8]. The association between FP-ERG and % L is nonlinear and as 100% is approached, very small changes in measured spectral sensitivities are associated with relatively large changes in estimates of % L. Thus, near 100%, small experimental errors led to some values that exceeded 100% upon normalization. These data combined with genetic analysis of the L and M cone genes to determine the spectral sensitivity of each opsin allowed for the determination of the L:M cone ratio.

#### Association study DNA preparation, pull-down, and sequencing

For each of these gene regions, we isolated DNA by targeting sequence intervals delineated by the end coordinate of the neighboring upstream gene to the start coordinate of the neighboring downstream gene using an oligo-based pull-down method (**Fig. S3**). If the neighboring upstream or downstream genes were less than 100 kb from the gene of interest, we expanded the pulled down region to include at least 100 kb of upstream and downstream sequence (**Fig. S3**). DNA was isolated from buccal swabs or whole blood using PureGene DNA extraction kits as described in [9]. DNA was then diluted to 10 ng/μL in low ETDA (USB corporation, #75793). DNA was sheared to an average length of 250 bps using the Bioruptor^®^ Pico (Diagenode). Samples were stored on ice for 10-15 min prior to sonication. Sonication was performed for 7 cycles of on for 15 seconds, off for 90 seconds. After 3 cycles, the samples were briefly centrifuged before continuing with sonication for the remaining 4 cycles. Library preparation and hybridization pull-downs were performed according to the SeqCap EZ HyperCap Workflow User’s Guide, version 2.0 following all recommended instructions. Libraries were created using the KAPA Hyper Prep Kit (KAPA Biosystems, #KK8504), using the KAPA Dual-Indexed Adapter Set (15 mM, KAPA Biosystems/Roche, #08 278 555 702). Probes for the pull-down were designed according to the Roche Nimble Design software, targeting the gene regions listed in (**Fig. S3**). Samples were sequenced on a Nova-seq, high-output, with 125 bp paired-end reads.

#### Association study analysis

Paired-end reads for 738 samples were aligned to the human genome (build hg38) using BWA mem (v0.7.15) with default parameters. Duplicate reads were marked using Picard (v2.9.2). Each region was subdivided into nonoverlapping 1 kb windows starting at the upstream boundary using bedtools (v2.29.2). Mapped reads were genotyped on a region-by-region basis using Freebayes-parallel (v1.2.0), the argument ‘--use-best-n-alleles 4’, and, for the region on chromosome X, ‘-p 1’. Variant calls were filtered for quality greater than 10 times the number of additional observations, at least one alternative allele read on each strand, and at least two reads supporting the alternate allele balanced up- and downstream using vcflib (v1.0.0). Duplicate variants were then removed. Variants were filtered with PLINK (v1.90b6.12) for a minor allele frequency of at least 3.5% and a per-variant genotyping rate of at least 90%. Variants were pruned using LDAK (v5.0) with a linkage disequilibrium cutoff of *R*^2^ < 0.75 within 100 kb windows. A kinship matrix was calculated from the pruned variant set using LDAK, ignoring weights and a power of -1. P-values were calculated in LDAK from the pruned variant set, using the first five genotype eigenvectors as covariates to control for potential population stratification. Significant variants were defined using an alpha of 0.05 after Bonferroni correction.

### QUANTIFICATION AND STATISTICAL ANALYSIS

#### HEK293 in situ hybridization quantification

Entire area of cell coverage was tile scanned and all individual cells inside the region were manually counted using the Adobe photoshop count tool. Cells were delineated by presence of pink, blue, or purple signal with a boundary drawn around the signal. Images were contrasted out to better determine brightfield boundaries if need be. Empty cells were not counted.

#### Human adult retina in situ hybridization quantification

For adult retinas, all serially sectioned punches of human retina were imaged and counted manually and identified by presence of pink, blue, or purple signal using the Adobe photoshop count tool. Exclusive expression of *M*-*opsin* or *L*-*opsin* was observed in cones. Co-expression of *M*-*opsin* and *L*-*opsin* was never observed in human retinas (purple signal). Cells were delineated by presence of pink, blue, or purple signal with a boundary drawn around the signal. Images were contrasted out to better determine brightfield boundaries if need be. Average ratios were calculated using a one-way ANOVA with Tukey’s multiple comparisons test. Graphs and statistical tests were done in GraphPad Prism version 9.3.1 for Mac, GraphPad Software, San Diego, California USA, www.graphpad.com. All error bars represent standard error of the mean (SEM).

#### Quantification of IHC and ISH BaseScope of M- and L-opsin stained HEK293 cells and adult fetal retina sections

IHC and ISH BaseScope images were manually aligned in Adobe photoshop using the hoescht nuclei stains. Images of the ISH BaseScope stain were used to count cells positive for *M*-*opsin* mRNA, *L*-*opsin* mRNA or both using the count tool. By overlaying the IHC stain, we determined which of these cells are also co-stained for M and L-opsin protein. Graphs and statistical tests were done in GraphPad Prism.

#### Human fetal retina in situ hybridization quantification

For the D122 human fetal retina, the entire eye was cryosectioned and representative sections were chosen, approximately 3 serial sections every 6 serial sections. Only *M*-*opsin* was observed in all sections with positive signal, which were all in the posterior pole of the eye.

#### Human opsin transcript pileups

Transcripts per million (TPM) values were then used to generate graphs in GraphPad Prism.

#### RNA sequencing graphs

All graphs and statistical tests were generated using GraphPad Prism.

#### Organoid quantification

All sectioned organoids were imaged and counted manually using the Adobe photoshop count tool. Organoids that had fewer than 150 cones (n ≤150) were removed from our analysis. A one-way ANOVA with Dunnett’s multiple comparisons test was used to determine significance. All error bars represent SEM.

#### Materials Availability

All reagents are available upon request. There are restrictions to the availability of cell lines listed in **Table 1** due to Material Transfer Agreement (MTA).

#### Data and code availability

Bulk organoid and Weri-Rb-1 RNA sequencing data are available upon request. Patients involved in the association study did not consent to include the release of raw sequencing or genotype data. The pipeline to analyze the M-versus L-opsin pileups is available at https://github.com/bbrener1/johnston_retina. The pipeline to analyze the association data is available at https://github.com/bxlab/2018.09.01_Eldred_GWAS/releases/tag/v1.0.

